# Development of a Bayesian network for probabilistic risk assessment of pesticides

**DOI:** 10.1101/2021.05.20.444913

**Authors:** Sophie Mentzel, Merete Grung, Knut Erik Tollefsen, Marianne Stenrød, Karina Petersen, S. Jannicke Moe

## Abstract

Conventional environmental risk assessment of chemicals is based on a calculated risk quotient, representing the ratio of exposure to effects of the chemical, in combination with assessment factors to account for uncertainty. Probabilistic risk assessment approaches can offer more transparency, by using probability distributions for exposure and/or effects to account for variability and uncertainty. In this study, a probabilistic approach using Bayesian network (BN) modelling is explored as an alternative to traditional risk calculation. BNs can serve as meta-models that link information from several sources and offer a transparent way of incorporating the required characterization of uncertainty for environmental risk assessment. To this end, a BN has been developed and parameterised for the pesticides azoxystrobin, metribuzin, and imidacloprid. We illustrate the development from deterministic (traditional) risk calculation, via intermediate versions, to fully probabilistic risk characterisation using azoxystrobin as an example. We also demonstrate seasonal risk calculation for the three pesticides.

## 1. Introduction

Pesticides play an important role in food production, by maintaining or enhancing crop yields and quality in arable farming. However, they can also lead to harmful effects in the environment and pose risks to human health. There is now a widespread concern about such regular emission of a substances designed to control specific target organisms and their effects on ecosystems (Van den Brink et al. (2018), Bradley et al. (2017), Mohaupt et al. (2020), Szöcs et al. (2017), Boye et al. (2019)). In spite of strict regulations of pesticide use (e.g. Directive 2009/128/EC; Regulation (EC) No 1107/2009), there are still knowledge gaps for potential environmental impact of these pesticides and their mixtures (Bradley et al. (2017), Szöcs et al. (2017), Mohaupt et al. (2020)). Current risk assessment methods use conservative assumptions to avoid underestimating the risk (Verdonck et al., 2003) and decision-makers rely on large safety margins for protective decision making (Fairbrother et al., 2015).

In general, risk assessment of pesticides is carried out to protect human health as well as the health and biodiversity of ecosystems (Schäfer et al., 2019). The purpose is to assess the probability that adverse effects of regulatory concern occur in ecosystems due to the exposure to one or several chemicals. This can be done as a prospective assessment for registration of substances before products enter the market, or as a retrospective assessment for potentially harmful substances that are already in use (Forbes & Calow, 2002). The environmental risk assessment process usually incorporates exposure and effect assessments as well as a risk characterization (Figure 1). Exposure assessment covers the estimation of predicted or measured environmental concentration (PEC) of the compound in the environment (van Leeuwen & Vermeire, 2007). PEC is usually calculated as the maximum environmental exposure concentration (Finizio & Villa, 2002). Effect assessment is typically based on the response of species that are exposed to a chemical in toxicity tests, such as data for toxicity endpoints (e.g. mortality, reproduction and growth) after short term (acute) or long term (chronic) exposure (van Leeuwen & Vermeire, 2007). Usually, a so-called predicted no-effect concentration (PNEC) is obtained from the most sensitive no-observed-effect concentration (NOEC). Alternatively, the PNEC can be calculated from the hazardous concentration for 5% of the species (HC5) based on the species sensitivity distribution (SSD) (Commission, 2003). To account for uncertainty, the lowest NOEC (alternatively the HC5) is divided by an assessment factor (AF) to derive the PNEC, so it can be considered a safe concentration for non-target organisms (Schäfer et al., 2019). Risk characterization includes a risk estimation by comparing effect (hazard identification and characterization) and exposure assessment, some of the metrics used are margin of exposure, hazard or risk quotient (Committee et al., 2019). To ensure low risk it is required that the PEC is lower than the PNEC (Commission (2003), Schäfer et al. (2019), so when using a risk quotient (RQ), it is derived by the ratio PEC/PNEC. If risk quotient exceeds 1 a risk of harmful effects to the environment is indicated.

**Figure 1.**
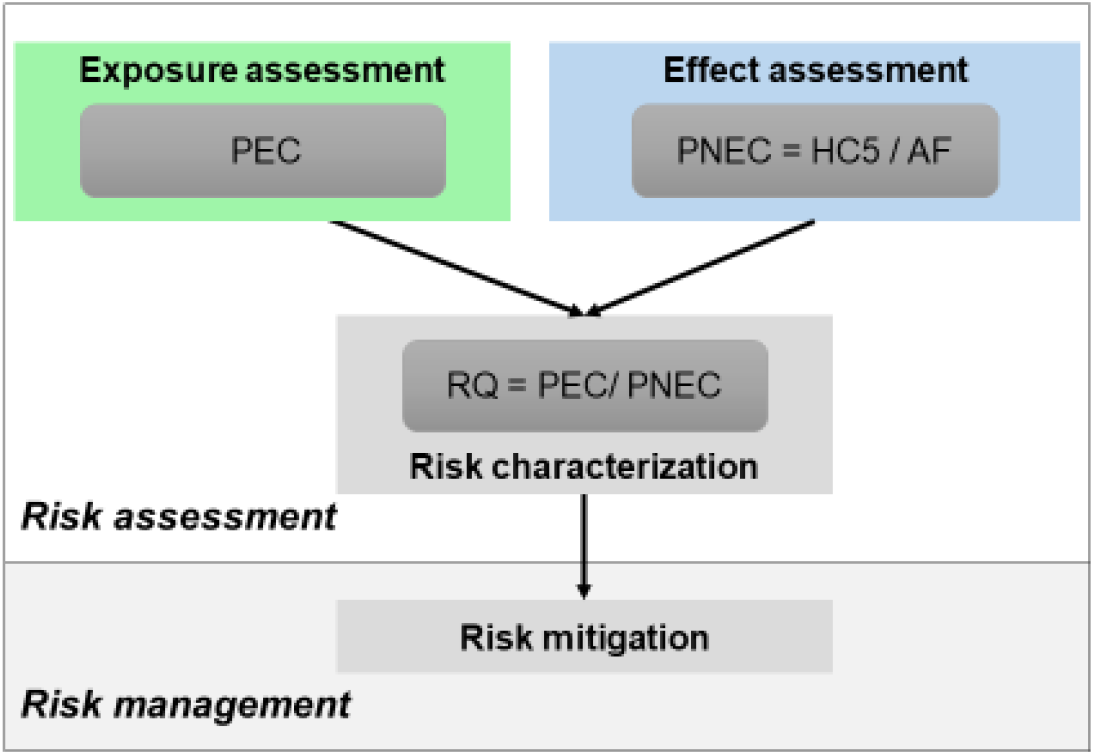
General ecological risk assessment process (RQ = Risk quotient, PEC = predicted or measured environmental concentration, PNEC = predicted no effect concentration, HC5 = hazardous concentration for 5% of the species derived from SSD (species sensitivity distribution), AF = assessment factor)

Risk is usually considered an estimation of the likelihood that an adverse effect occurs on a biological target when being exposed to a chemical (Finizio and Villa (2002), S Jannicke Moe et al. (2021), Fairbrother et al. (2015)). Nevertheless, in the commonly used framework for environmental risk assessment, the output of risk characterisation tends to be single value (the risk quotient) from which the conclusion is a “yes/no” statement (Fairbrother et al., 2015). It has been argued that such single-value estimates cannot stand alone as a scientifically defensible characterization of ecological risk (Campbell et al., 2000). The analysis and quantification of uncertainty is a vital part of risk assessment of environmental impacts of pesticides, which is not reflected in the single-value risk estimate (USEPA (2014), Fairbrother et al. (2015)). Based on this, a concerted action was established to develop a European framework for probabilistic risk assessment of the environmental impacts of pesticides (EUFRAM). The consortium named several shortcomings of conventional ERA (EUFRAM 2006), for example: there is no indication of the level of certainty associated with the risk assessment; no quantification of the risk is carried out; the uncertainty calculation is not transparent but hidden in assessment factors; and it is difficult to follow all steps of the risk assessment. Various recommendations were given for development towards probabilistic risk assessment, mainly based on the use of cumulative probability distributions (EUFRAM, 2006). Nevertheless, non-probabilistic methods are still more commonly used (Fairbrother et al., 2015). One reason can be a lack of training in probabilistic methods and tools in ecotoxicology.

The objective of this study was to explore Bayesian network modelling as a tool to combine probability distributions of pesticide exposure and effects, to facilitate the calculation of the risk quotient as a probability distribution instead of a single number. We aimed to align the developed model to the European Union (EU) regulatory requirements and current risk assessment procedures, to enable comparison of the Bayesian network approach with the existing approaches. To this end, we present the development from a deterministic towards a fully probabilistic Bayesian network approach to risk characterisation. The model application is demonstrated for three examples of pesticides and for different seasons.

## 2. Approaches to probabilistic risk assessment

### 2.1 Proposed methods for probabilistic risk assessment

Probabilistic risk assessment has been defined as using “*probabilities or probability distributions to quantify one or more sources of variability and/or uncertainty in exposure and/or effects and the resulting risk*” (EUFRAM, 2006). This allows the inclusion of estimates of uncertainty and stochastic properties (Solomon et al., 2000). There are now several probabilistic methods in use for risk characterisation. The species sensitivity distribution (SSD) (Posthuma et al., 2001) is a probabilistic model for the variation in sensitivity of biological species to a single or a set of toxicants, which is used in several frameworks (Belanger & Carr, 2020). Guidance on modelling and data requirements can be found in the “Technical Guidance for Deriving Environmental Quality Standards” (SCHEER, 2017). Many of the probabilistic methods currently at hand also incorporate a distribution for the exposure part. An overview probabilistic methods currently at hand is given **Error! Reference source not found**.. Methods such as quantitative overlap and joint probability curves are relatively easy to construct (Verdonck et al. (2003), Campbell et al. (2000)), and use more available data for exposure and effect compared to traditional approaches (Campbell et al., 2000). They also allow for an estimation of likelihood of potential ecosystem impact and their magnitude (Solomon et al., 1996). Recently, an “Ecotoxicity Risk Calculator” was presented by Dreier et al. (2020) that uses joint probability curves. It is able to express more information than a single value risk quotient, as it depicts the relationship between cumulative probability and magnitude of effect. The use of both effect and exposure distributions enables a more powerful approach for risk assessment and communication (Dreier et al., 2020). However, most of these methods do not provide exact quantifications of magnitudes and likelihoods of potential effects, they do not make quantitative predictions and only estimate relative risks (Solomon et al. (2000), Hall et al. (2000)), which can be hard for decision-makers to understand and interpret (Verdonck et al., 2003).

### 2.2 From deterministic to probabilistic risk quotient

Another method more consistent with the probabilistic definition of risk is the calculation of probabilistic risk quotients. It can be useful for ranking of different scenarios as well as prioritizing among alternative risk scenarios (Campbell et al., 2000). A fully probabilistic risk quotient calculation requires the quantification of a probability distribution for both exposure and effect. In cases where exposure or effect data are too limited, an alternative “intermediate” probabilistic approach could be applied by using a distribution for either the exposure or effect component (Figure 1). This will allow for some variability to be taken into account when deriving a distribution for the risk quotient. For example, an intermediate approach could be applied when an effect concentration distribution can be quantified by a species sensitivity distribution, although few exposure measurements are available. Figure 2 displays the underlying concepts of the traditional deterministic approach and the intermediate and fully probabilistic approaches. The traditional deterministic approach (Figure 2a) used single-value PEC and PNEC single value risk quotient. The second option (Figure 2b) used an exposure distribution together with a single value PNEC, derived the same way as in the traditional approach. Though, unlike the traditional approach, here a risk quotient distribution is derived. The third option (Figure 2c) uses the probability distribution of effects (corresponding to an SSD). Instead of using the SSD to extract a single-value HC5 as a basis for a single-value PNEC in combination with an assessment factor, in this case, an uncertainty factor (UF) is applied to the calculated exposure/effect ratio distribution. The uncertainty factor plays a similar role as an assessment factor, that is to adjust the predicted risk to account for uncertainties e.g. associated with extrapolation from laboratory toxicity tests to environmental effects. However, we chose to use the slightly different term “uncertainty factor” to avoid misusing the more well-established term “assessment factor”. For the fourth option (Figure 2d), probability distributions are calculated for both exposure and effect distributions. Again, no PNEC is derived, so after calculating the exposure/effect ratio distribution, an uncertainty factor is applied to adjust the risk quotient distribution.

**Figure 2.**
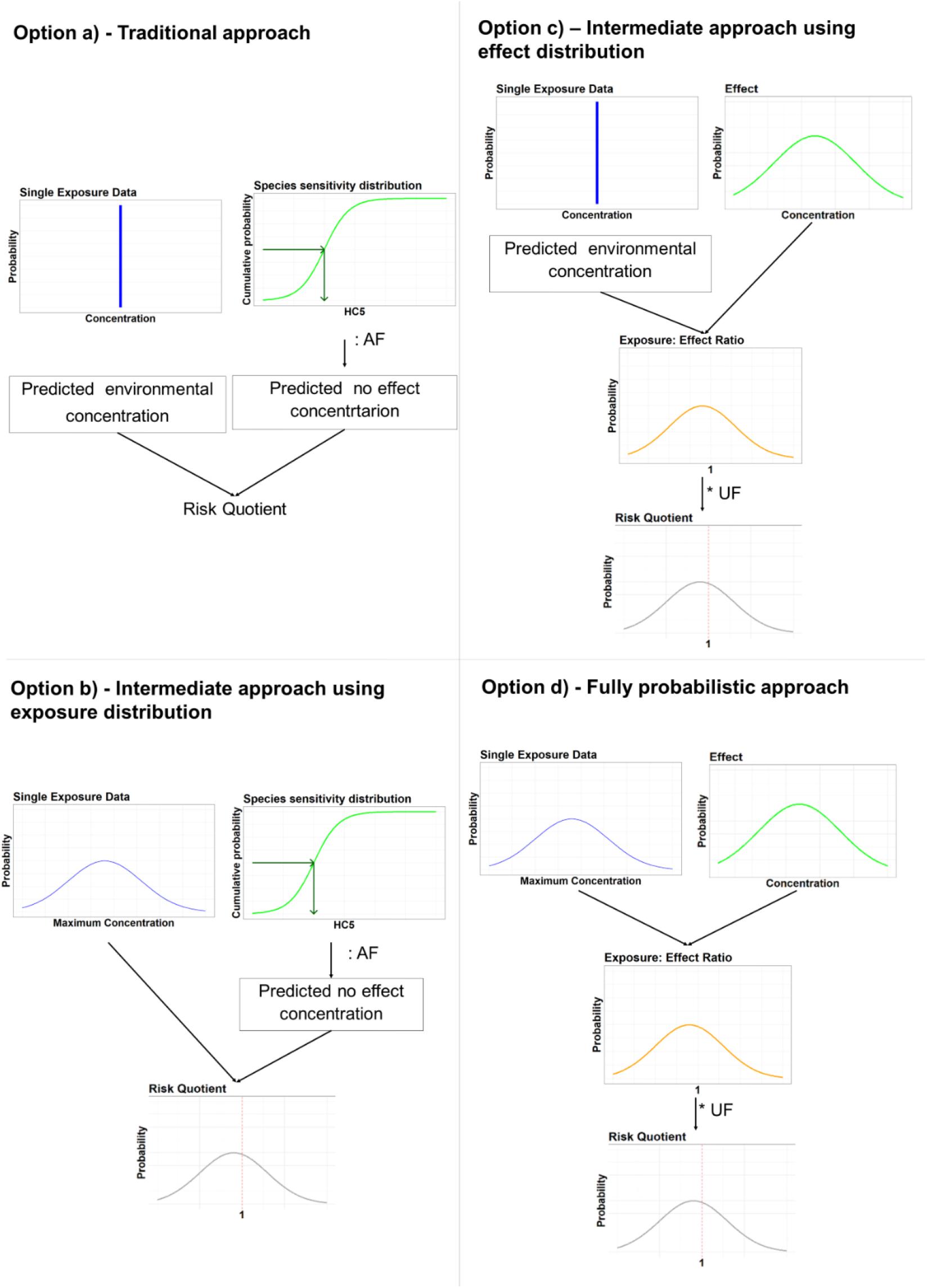
Systematic overview of the traditional approach to derive a risk quotient, compared to two intermediate probabilistic options and a fully probabilistic option that derive a risk quotient distribution.

### 2.3 Probabilistic risk assessment using Bayesian networks

The early efforts of probabilistic risk assessment for pesticides, which were usually visualised by cumulative distribution curves, were sometimes difficult to interpret for both for advanced users and the general public (EUFRAM 2006). As an alternative, Bayesian networks may provide a way to overcome limitations associated with visualization of risk estimations while accounting for uncertainties when using probabilistic approaches. They have been recognized as a tool to analyse complex environmental problems and support decision making while considering uncertainty (Sperotto et al., 2017), and have lately been increasingly used for environmental risk assessments (Moe et al. 2021). A Bayesian network can characterize a system by showing its interactions between variables in a network (Chen & Pollino, 2012) through a directed acyclic graph (Kanes et al., 2017). They are probabilistic graphical models implementing Bayes’ rule for updating probability distributions based on evidence. The nodes (variables) have discrete states (e.g. intervals), quantified by discrete probability distributions. The causal links (arrows) represent conditional probability tables (CPT) which can be based on equations. The degree of belief (probability) that a variable will be in a particular state given the state of the parent variables, as specified by the conditional probability table (Chen & Pollino, 2012), and by using Bayes’ rule for updating probability distributions based on new evidence (Molina et al., 2010). In this project, Bayesian network construction followed guidelines provided by Marcot et al. (2006) and Pollino and Henderson (2010).

Bayesian networks have an integral feature suitable for risk estimation as they present results in probability distribution form instead of point estimates. They can accommodate different kind of data; its sources can include both direct measurements and output from models. Also, if data are limited or non-existent, it is possible to include expert opinions instead (Pitchforth & Mengersen, 2013). The models can be updated with new information on pesticide exposure and effects whenever it becomes available. Model updates are carried out by combining prior probabilities and new data so that an update of the network posterior probabilities can take place as a response to the added observational information (Franco et al., 2016). Bayesian networks are especially useful for pesticide risk assessment and management tasks as these require characterisation of the uncertainties (Carriger and Newman (2012)). Focusing on a terrestrial species (puma), Carriger & Barron (2020) displayed a process of mapping cause-effect relations into a quantitative model. This is supported by *Catenacci & Giupponi (2013)* who found that the Bayesian network approach can examine different phenomena due to its flexibility for interdisciplinary integration, e.g. climatic, physical, ecological, and socio-economic (Catenacci & Giupponi, 2013). They also have the ability to perform predictive (forward), diagnostic (backward), and mixed (forward and backward) inference (Carriger & Barron, 2020).

## 3. Methods

### 3.1 Study area

The model was developed based on monitoring data from a catchment within the Norwegian Agricultural Environmental Monitoring Program (JOVA) located in South-East Norway (Heia, location: 59°21’29’’N, 10°47’52’’E). The monitoring catchment has a total area of 1,7 km^2^ of which 62% are cropland. As the catchment is located in a coastal climate, winters are mild and the growing season starts relatively early as compared to Norwegian conditions in general. The catchment has an annual rainfall of 829 mm and a mean annual temperature of 5.6 °C (in 2016). The crop production in the catchment is mostly grain (up to 75%). Potato and vegetable production made up about 40% until 2007 and had decreased to about 25% in 2015. The catchment’s use of plant protection products and exposure data are recorded in the JOVA program (Bechmann et al., 2017). Flow-proportional composite sampling of stream water at the catchment outlet was performed in the JOVA program throughout the spraying season and the analysis of concentrations of a wide range of current and previously used pesticides were included. Based on these data, exceedance of environmental safety thresholds are identified for different agricultural management practices for key agricultural production systems in various catchments in Norway (Stenrød, 2015). The JOVA monitoring data for pesticides has been collected through 25 years (1995 onwards) and thus also support the retrospective assessment of ecological risk and temporal trends (Bechmann et al., 2017).

### 3.2 Pesticides - exposure and effect data

The chemicals selected for analysis in this study are most frequently occurring pesticides and highest in concentration in the study catchment (Table 1). Azoxystrobin and metribuzin are approved chemicals for use in the EU and Norway. Since 2013 the use and sale of Imidacloprid is prohibited in the EU (Commission, 2013). Of the selected chemicals, only the fungicide azoxystrobin has low solubility in water at 20 °C (6.7 mg L^-1^), whereas metribuzin and imidacloprid have high solubility in water. All pesticides form metabolites primarily in soil.

**Table 1.**
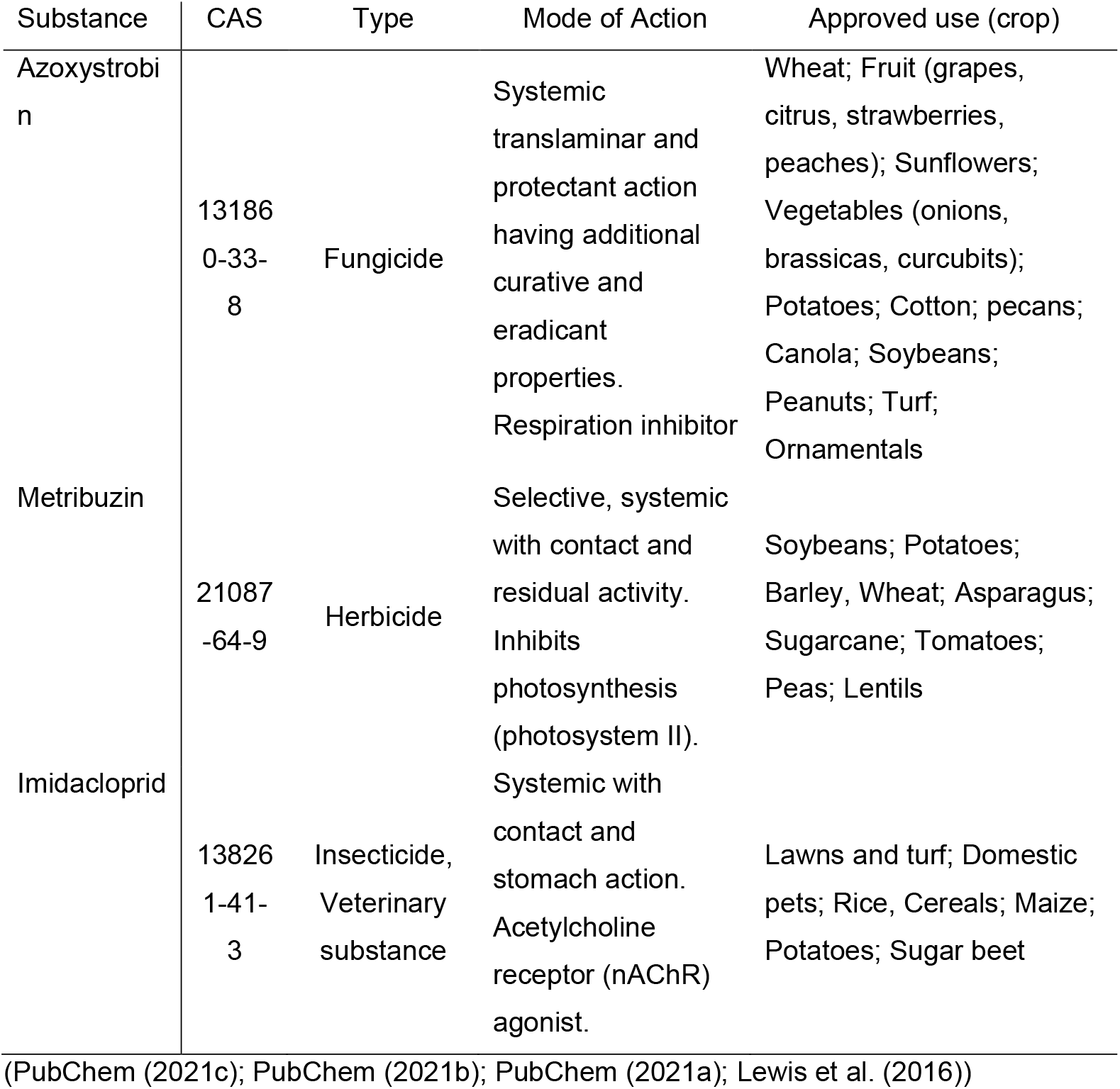
Information about selected pesticides their Chemical Abstract Service (CAS), pesticide type, mode of action and common application crop.

The data used in this study were obtained from the NIVA Risk Assessment database (NIVA RAdb, www.niva.no/radb), which hosts exposure and effect data from a wide variety of sources. Moreover, this database provides transparent and harmonized cumulative risk predictions according to international recommendations for harmonised approaches for human and ecological risk assessment (Tollefsen, 2021). Exposure data for the period 11.05.2011 to 06.12.2016 from the JOVA monitoring program were extracted from NIVA RAdb database.

The total number of measured environmental concentrations was 55 for azoxystrobin, and 59 for metribuzin and imidacloprid. There is large variation in the measured concentration levels during the season and years for each of the pesticides. The percentage measurements below the limit of quantification (LOQ) were 53%, 16, % and 11% for azoxystrobin, metribuzin and imidacloprid respectively. In general, sampling of pesticides varied greatly between the years and month with higher concentrations in summer and autumn and lower concentrations in spring and winter.

For the selected pesticides, toxic effects data for several freshwater species representing various taxonomic groups were extracted from the NIVA RAdb. The data set consisted of NOECs (no observed effect concentration) for adverse effects such as growth, reproduction, and population. For each chemical, multiple NOEC values from the same species were used in our analysis (see Table 2). In traditional effect assessment, only the most sensitive value per species is often chosen to derive an SSD, although in some cases an average is also used.

**Table 2.**
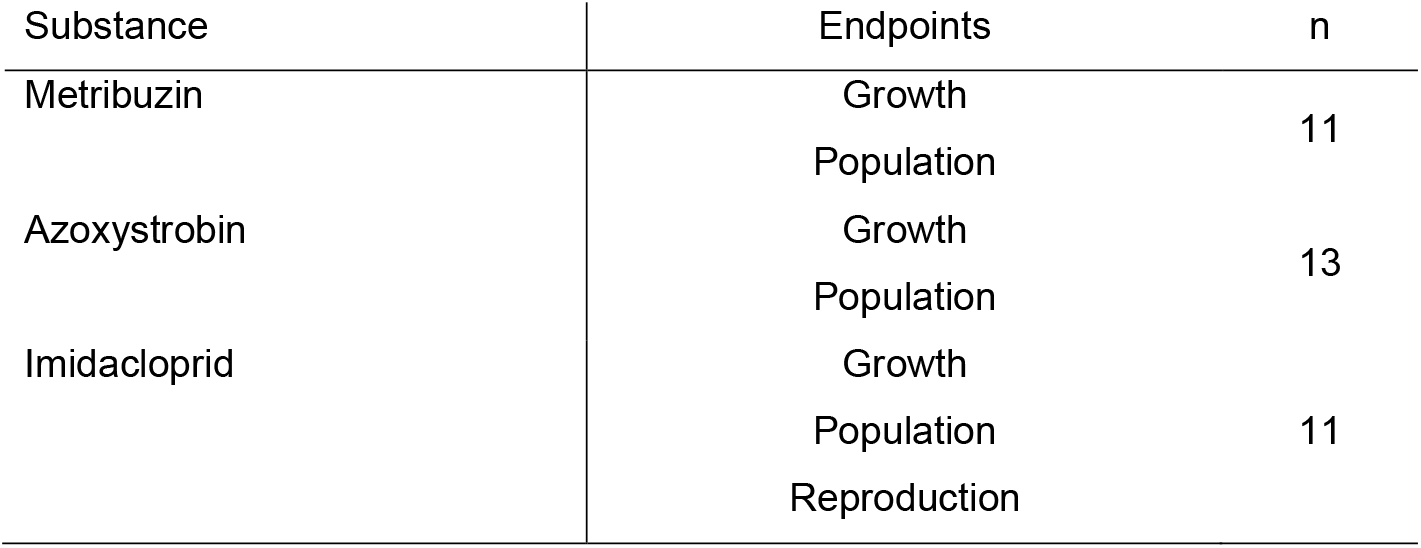
Overview of collected effect/ toxicity data for the selected pesticides, also showing their adverse effect endpoint, n = number of means used to fit the distribution and species with multiple NOECs for the same substance

| Substance | Endpoints | n |
| --- | --- | --- |
| Metribuzin | Growth | 11 |
|  | Population |  |
| Azoxystrobin | Growth | 13 |
|  | Population |  |
| Imidacloprid | Growth | 11 |
|  | Population |  |
|  | Reproduction |  |

### 3.3 Data Processing

Data preparation was carried out with R version 4.0.2 (Team, 2020) using packages including *tidyverse* (version 1.3.0) (Wickham et al., 2019), *dplyr* (version 1.0.2) (Wickham et al., 2020) and *readxl* (version 1.3.1) (Wickham & Bryan, 2019). To obtain probability distributions for the BN model from the exposure and effects data, log-normal distribution models were fitted to the data using the *R* package *MASS* (version 7.3-51.6) (Venables & Ripley, 2002).

In the case of exposure data below Limit of Quantification (LOQ), new values in the range from 0 to LOQ were simulated using mean and standard deviation from the fitted log-normal distribution. To take into account the seasonal variation in pesticide exposure, a separate probability distribution was estimated for each season, defined as follows: Winter = Dec-Feb; Spring = Mar-May; Summer = Jun-Aug; Autumn = Sep-Nov.

For the effect distribution, likewise, a log-normal distribution was fitted to the NOEC values available for each pesticide. In cases where multiple NOEC values of the same species were present, the mean NOEC was used. The fitted distribution corresponds to a species sensitivity distribution (SDD), which is often fitted as a log-normal distribution (Belanger & Carr, 2020). However, while SSDs are traditionally used to derive a single PNEC value (Figure 1), we used the whole probability distribution of effects data in this study. For comparison with the traditional risk quotient calculation based on a PNEC, as described in introduction a HC5 was derived from a species sensitivity distribution using the package *ssdtools* (Thorley & Schwarz, 2018).

### 3.4 Parameterization of the Bayesian networks

The Bayesian networks were built in Netica (Norsys Software Corp., www.norsys.com). For each pesticide, a BN was built with identical structure except for the range the exposure and effect concentrations were discretized. The individual node description can be found in Table 3; further detailed information can be found in the Supplementary material.

**Table 3.** Node description for the example of Option d the fully probabilistic approaches (see Figure7d), also describing the discretization type, number of states, conditional probability table input and parent relation

| Node / Variable | Type of discretisation | States |
| --- | --- | --- |
| Exposure concentration distribution | C | 10 |
| Effect concentration distribution | C | 10 |
| Exposure - effect - ratio distribution | C | 8 |
| Uncertainty factor | D | 7 |
| Risk quotient distribution | C | 8 |

\* D: Discretized discrete
\*\* C: discretized continuous; continuous variables were binned into the states
\*\*\* States: number of intervals of each node
\*\*\*\* further details about the Node input and assumption see Supplement Information

For both exposure and effects nodes, the range was defined by the observed values of the given pesticide, and the intervals were discretized into 12 equidistant bins in log10-scale. The fitted log-normal distributions were used to parameterize the parent nodes (*for more information about input and equations used see Supplementary material*). The probability distribution of the nodes “Exposure Concentration (µg/L)” and “Effects Concentration (µg/L)” was calculated from their respective parent nodes by 10exp-transformation. The node “Exposure/Effect Ratio” was discretized in 8 equidistant bins and calculated by the equation [Exposure Concentration (µg/L)]/ [Effects Concentration (µg/L)]. Thereafter, the risk quotient distribution was derived by multiplying the “Exposure/ Effect Ratio” with an uncertainty factor. The uncertainty factor can be applied to account for uncertainties in the effect assessment, similar to the use of an assessment factor in traditional risk assessment (Figure 1).This factor can be transparent and standardized in a simple manner by considering the information used during the effect assessment e.g. number of data points (Figure 3). In our model (Figure 1), the node “Uncertainty factor” have alternative levels that can be selected by the risk assessor, depending on the sources of uncertainty to be accounted for in the risk assessment.

**Figure 3.**
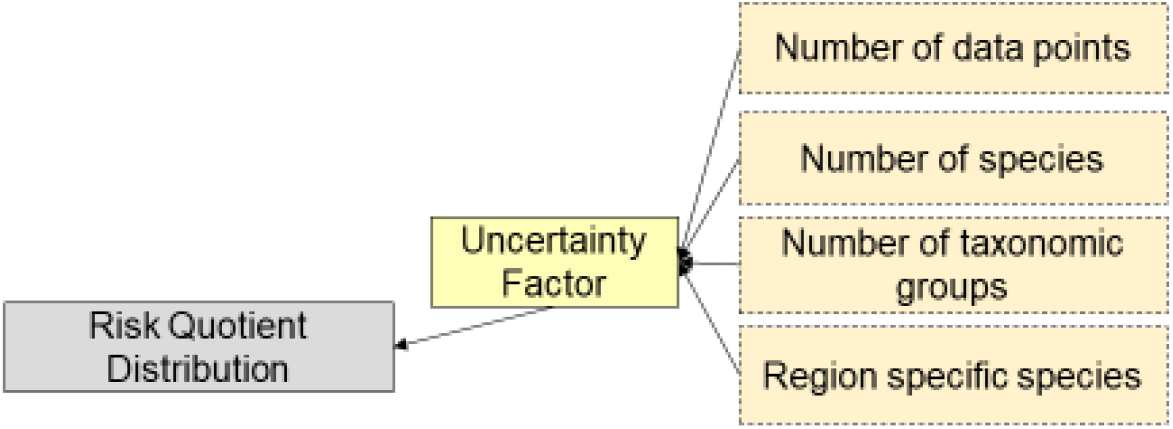
Possible sources of uncertainty that can be incorporated in the uncertainty factor

After the Bayesian network was constructed and parameterized a sensitivity analysis was carried out in Netica (Norsys Software Corp., www.norsys.com). One of the benefits of using this software is the simple execution of sensitivity analysis that can easily selected from the menu bar. The report displayed that the risk quotient distribution is dominated by the exposure side over the effect side, which is most likely due to the wider range of concentrations.

This way, a Bayesian network model is intended as a tool for calculating the risk quotient as a probability distribution, to account for e.g. temporal variability in exposure, taxonomic variability in effects, and other types of uncertainty.

## 4 Results and Discussion

### 4.1 Input values, distributions and uncertainty factor used of the Bayesian network

This section describes the parameterised version of the Bayesian network for each of the three pesticides, illustrated with azoxystrobin as an example. For comparison, the risk quotient was also calculated by the traditional single-values method (Figure 2a) as well as by the two intermediate options (Figure 2b and c). For the single-value exposure versions (Options a and c), the minimum (0.01 ug/L), mean (0.063 ug/L) and maximum (0.660 ug/L) of the measured concentrations were selected as alternative PEC values. The highest exposure concentration is usually used as the more conservative or protective choice. For the single-value effect version (Options a and b), the PNEC values were derived from an HC5 of 3.87 µg/L divided by an assessment factor of 10, 5, 3 and 1 (Figure 4).

**Figure 4.**
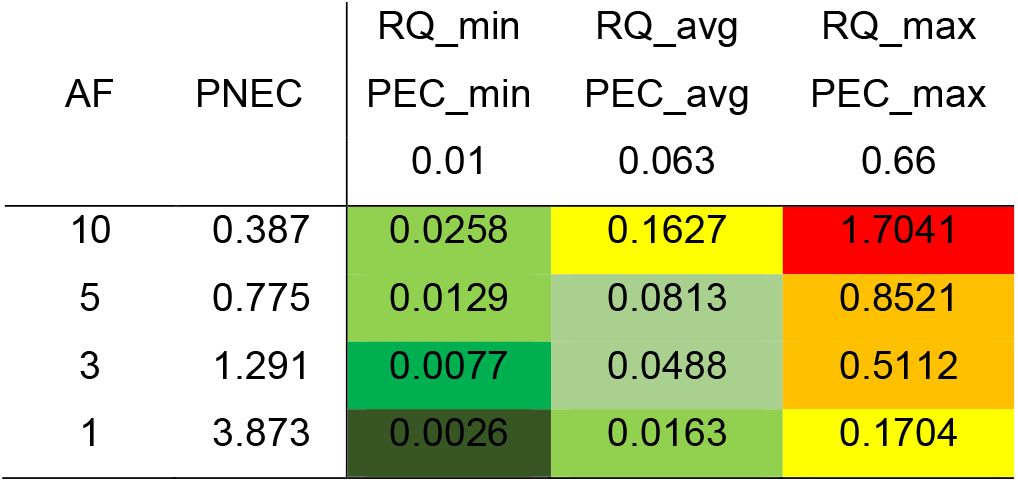
Risk quotient derived for minimum, average and maximum PEC and a PNEC (for Assessment factor of 1, 3, 5, and 10)

The probability distributions of exposure and/or effects data in Options b, c and d were based on the fitted log-normal distribution with mean and standard deviation. The exposure distribution had a mean of -4.148 ln (ug/L) with a standard deviation of 1.484 ln (ug/L). The effect distribution had a mean of 2.322 ln (ug/L) with a standard deviation of 0.56 ln (ug/L).

The seasonal version of the Bayesian network was parameterized with exposure distributions based on seasonal mean values for the three pesticides. Winter season had too few measured environmental concentrations to derive a distribution for all three chemicals and was therefore excluded from further analysis. In general, mean concentration in summer were higher than in spring and intermediate in autumn (Table 4). Except from Imidacloprid which has higher concentrations in autumn.

**Table 4.** Estimated mean and standard deviation of the exposure by season and effect distributions, which are used as input for the nodes in the Bayesian network.

| Compound |  | Exposure |  |  | Effect |
| --- | --- | --- | --- | --- | --- |
|  |  | Spring<br>ln (ug/L) | Summer<br>ln (ug/L) | Autumn<br>ln (ug/L) | ln (ug/L) |
| Azoxystrobin | mean | -5.029 | -3.939 | -4.018 | 2.322 |
|  | sd | 0.712 | 1.529 | 1.541 | 0.568 |
| Metribuzin | mean | -4.357 | -2.794 | -3.292 | 4.946 |
|  | sd | 0.966 | 1.416 | 1.363 | 2.432 |
| Imidacloprid | mean | -3.902 | -3.404 | -1.783 | 6.484 |
|  | sd | 1.481 | 1.116 | 1.743 | 4.004 |

The uncertainty factor was derived by diagnostic inference by instantiating the nodes for exposure, effect and risk quotient (Figure 5). For the exposure and effect concentrations, the intervals were set according to the mean of the observed values.

**Figure 5.**
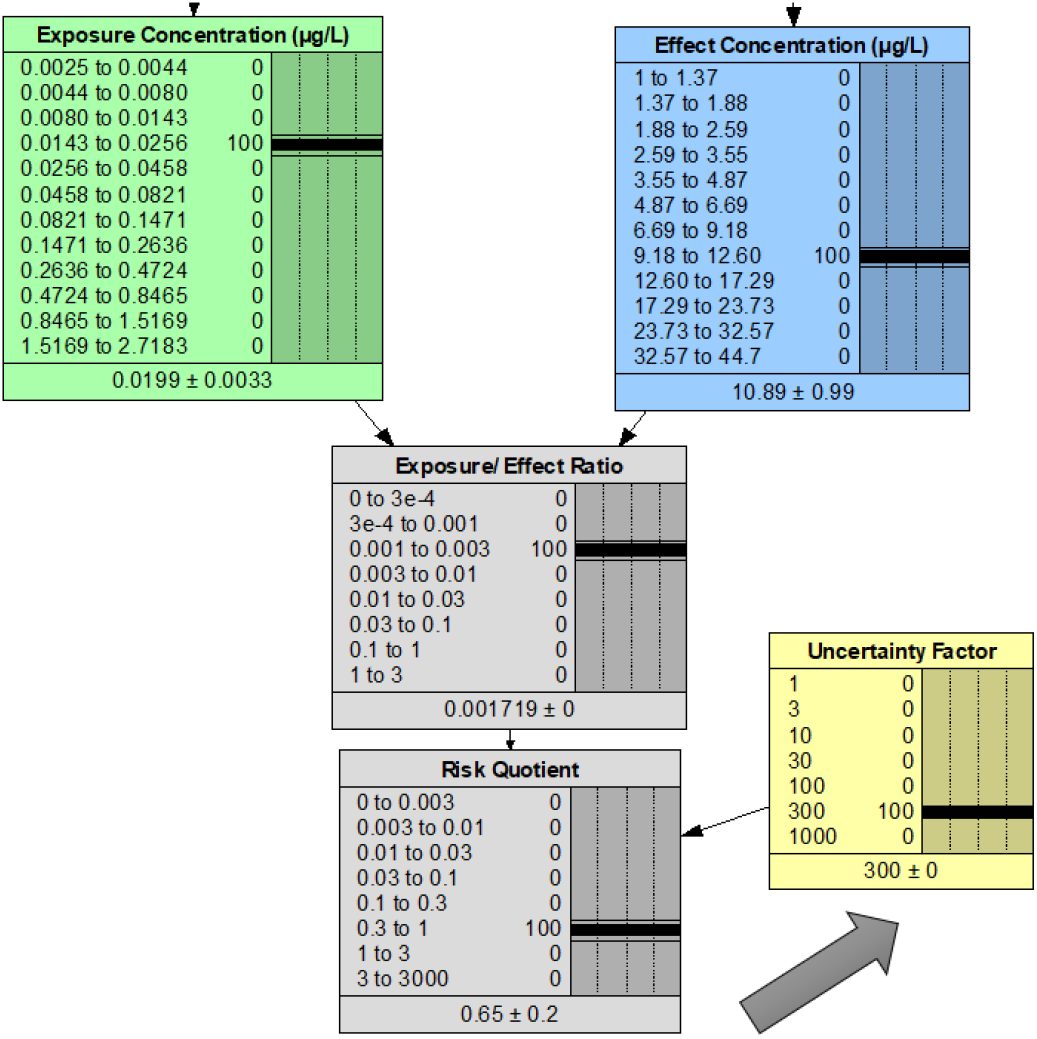
Example of diagnostic inference for this case study for a mean exposure and effect interval.

The appropriate uncertainty factors found corresponding to the assessment factors are displayed in the following Table 5. We chose uncertainty factors of 10, 30 and a 100 for the first example with Azoxystrobin and an uncertainty factor of 100 for all the seasonal versions of the Bayesian network.

**Table 5.** Uncertainty factors corresponding to assessment factors

| AF | PEC_min | PEC_avg | PEC_max |
| --- | --- | --- | --- |
|  | 0.01 | 0.063 | 0.66 |
| 10 | 10 | **100** | 1000 |
| 5 | 10 | **30** | 300 |
| 3 | 3 | 30 | 300 |
| 1 | 1 | **10** | 100 |

### 4.2 Risk quotient distributions predicted by the Bayesian network

The Bayesian networks for the different options for the risk quotient calculation were carried out for azoxystrobin and are displayed in Figure 6. For the Bayesian network approach, the risk quotient distribution node output was displayed for the different events and node settings. The colours range from green (no risk) to red (posing a risk) (Figure 7). The risk quotient distribution for the approaches ranged from 0 to 3000. Higher assessment factor and uncertainty factor can lead to the risk quotient > 1. The calculated risk quotients can be found in in Figure 4. An example using a BN approach for Option a, is displayed in Figure 6a. In this example the risk quotient was calculated using a mean PEC and a PNEC with an applied assessment factors of 5 and 10. The risk quotient distribution is estimated to be within the interval “0.03 to 0.1”.

**Figure 6.**
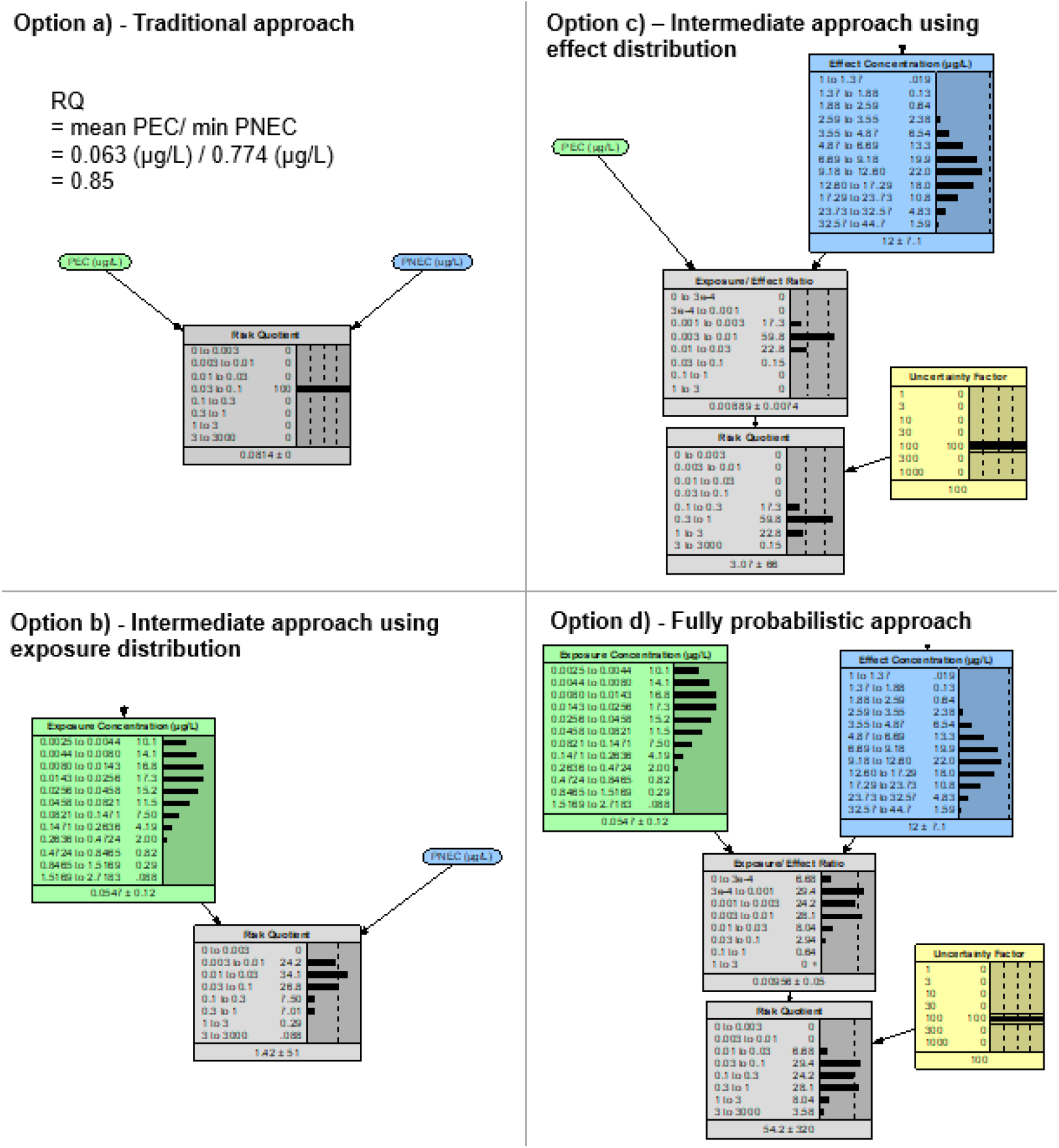
Example of Bayesian network for both intermediate and the fully probabilistic approach for the fungicide azoxystrobin, b) risk quotient distribution is derived for the PNEC derived with an Assessment factor of 5, c) for a mean PEC and uncertainty factor of 100, and d) distributed exposure and effect concentration, and uncertainty factor of 100.

**Figure 7.**
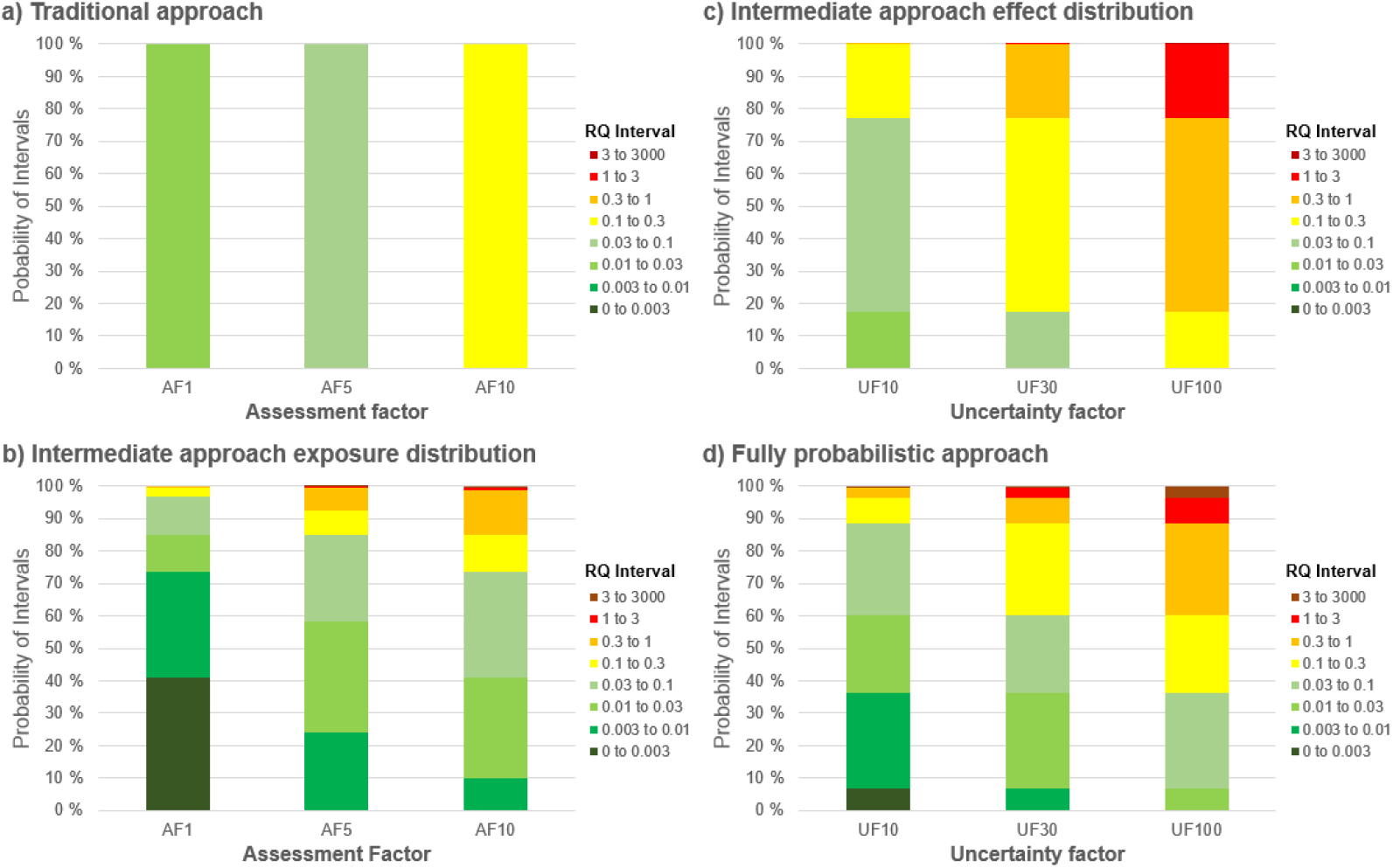
Risk quotient derived from the traditional approach using single mean PEC and PNEC values (a), and risk quotient distribution output from the Bayesian network for intermediate approaches with exposure distribution and PNEC (b), and mean PEC and effect distribution, with uncertainty factor 10, 30 and 100 (c), and a fully probabilistic approach with exposure and effect distribution and uncertainty factors 10, 30 and 100 (d).

When using an assessment factor of 1 and 10 the probability for the risk quotient to be in the interval of “0.01 to 0.03” and “0.1 o 0.3” is 100% (Figure 7a). Option b uses an exposure distribution and the same assessment factors as in Option a to derive the risk quotient, which is distributed over the intervals “0 to 0.0003” and “1 to 3”. For an assessment factor of 1 the probability for the risk quotient to be in an interval higher than 0.1 is about 3.2 % whereas for an assessment factor of 5 it is 26.4%. Option c in this example uses uncertainty factors calculated in Table 5. For the events of a mean PEC with an uncertainty factor of 100 the interval of “0.03 to 1” has the highest probability. If a uncertainty factor of 30 is chosen the interval of “0.1 to 0.3” instead has the highest probability (Figure 7c). The probability for the risk quotient to be above 0.3 with an uncertainty factor of 10 is less than 1%, with one of 30 it is about 23% and with one of 100 it is about 83%. The fully probabilistic approach – Option d uses distributions for both exposure and effect, when using an uncertainty factor of 10, 30 and 100. The probability for the risk quotient to be above 0.3 is about 4% with an uncertainty factor of 10, 12 % with one of 30 and about 40 with one of 100 (Figure 7d).

### 4.3 Seasonal variation in risk quotients

A more temporally refined version of the Bayesian network is displayed for the compound azoxystrobin (Figure 8), and used for calculating seasonal risk quotients for all three pesticides. The uncertainty factor was set to 100 as this was found to be most appropriate in comparison with the deterministic method Table 5. According to this model (Figure 8), the probability of the risk quotient for azoxystrobin exceeding 0.1 during summer is about 72%, while the probability of risk quotient exceeding 1 is about 15%.

**Figure 8.**
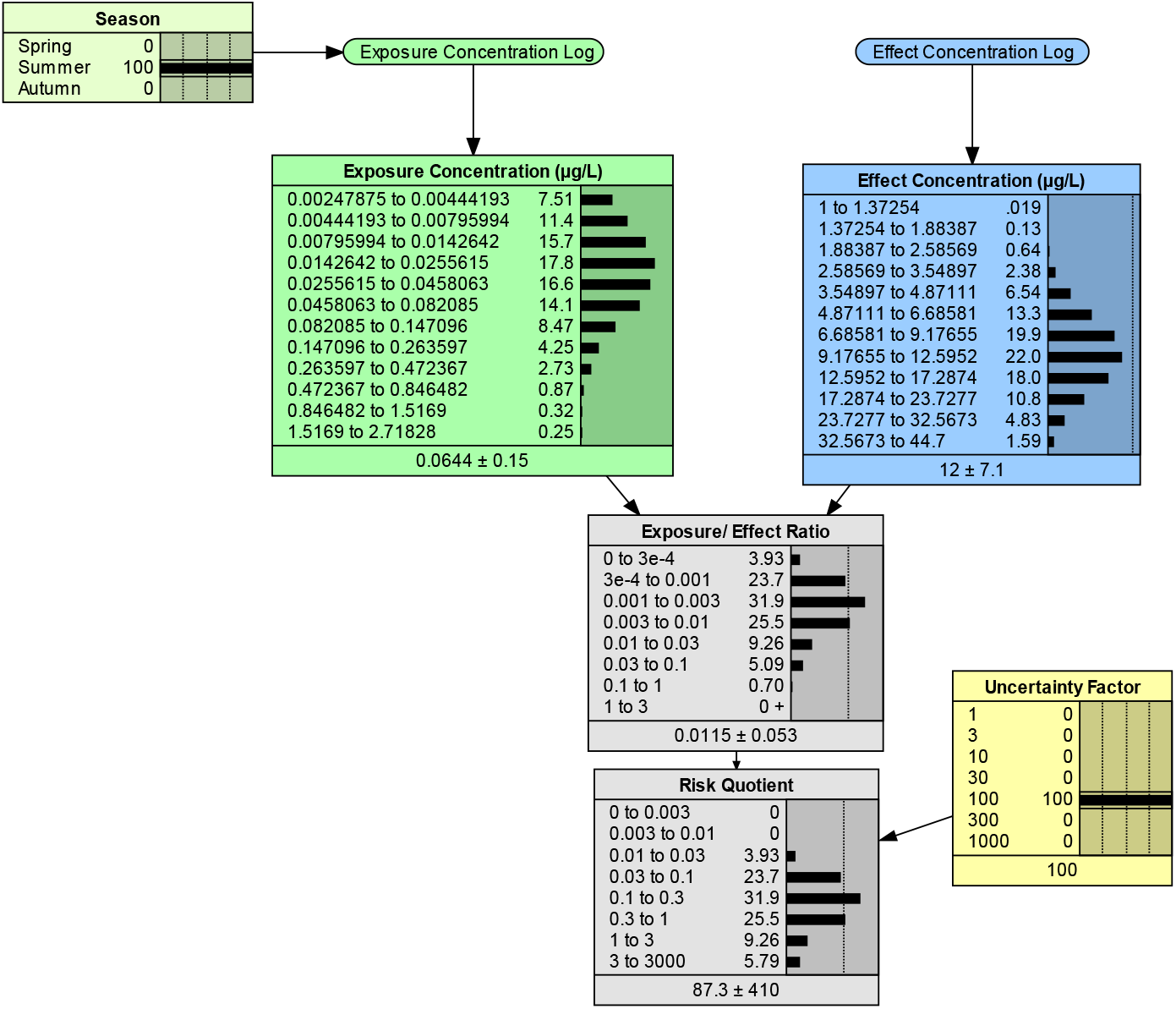
Example of a seasonal version of the Bayesian network model parameterised for the fungicide azoxystrobin, with application of an uncertainty factors of 100 for summer season.

In comparison with the two other pesticides, azoxystrobin clearly imposed a higher probability of exceeding the risk quotient levels of 0.1 to 0.3, especially in summer and autumn (Figure 9). Metribuzin and imidacloprid have a wider distribution for the risk quotient, mainly ranging from 0.0001 to 0.001. Spring and autumn distribution of probability in the case of imidacloprid are more similar, unlike azoxystrobin and metribuzin where summer and autumn appear to be more similar. These two seasons have higher probabilities for the risk quotient to be between above 1 than the spring season.

**Figure 9.**
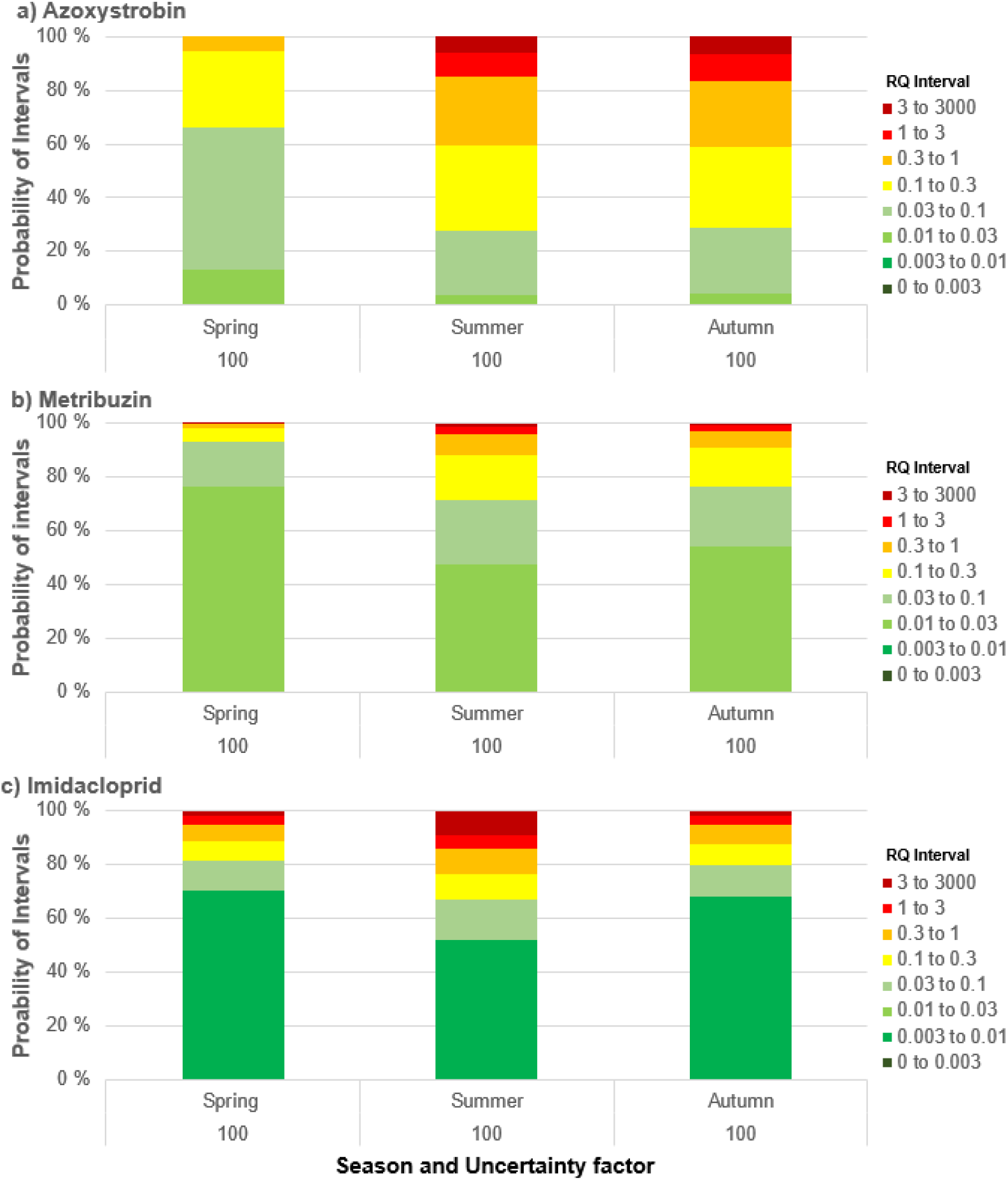
Calculated probability distribution of risk quotient, for spring, summer, autumn and uncertainty factors 10 for a) azoxystrobin, b) metribuzin and c) imidacloprid.

### 4.4 Evaluation of the Bayesian networks approach for risk characterisation

This study has demonstrated that Bayesian networks can account for quantified uncertainties and variabilities in a more coherent and transparent way than traditional risk characterisation. When developing this Bayesian network approach, we aimed at following important recommendations for probabilistic risk estimation described by EUFRAM (2006). We tried to accomplish these by accompanying the new methods with the conventional “deterministic” assessment, to enable that end-user (e.g. regulators) can become acquainted with the new methodology. Furthermore, the developed models follow well-known concepts described in the Technical Guidance Document (TGD) for whenever it was possible and logical. The TGD for example describes what an appropriate assessment factor is depending on the available data and mentions requirements for the used data for minimum amount of taxonomic and species used for SSD modelling (Committee et al., 2019). In addition, we tried to display the results in bar plots instead of cumulative probability. This was also pointed out by EUFRAM (2006) which mentioned stakeholders being more likely to take up results if they and the concepts used are as simple a possible and aligned with existing frameworks (EUFRAM, 2006).

Bayesian networks are increasingly used in environmental risk assessment (S. J. Moe et al., 2021). They can offer a transparent way of evaluating the required characterization of uncertainty for pesticide risk assessment as well as for ecological risk assessment in general (Carriger & Newman, 2012). Moreover, their application is not only carried out for risk estimation (e.g. risk quotient) it is also used to predict ecological effect more directly (e.g. decline in species abundance (Mitchell et al., 2021). *Dreier et al. (2020)* pointed out that the use of effect and exposure distribution allow for a competent risk assessment and communication approach. In their “ecotoxicity risk calculator”, they used joint probability curves/ risk curve based approach that is able to show the connection between cumulative probability and magnitude of effect (Dreier et al., 2020). Although this might be an advantage of using joint probability curves, probabilistic risk quotients can give a better sense of the risk estimates and are useful for ranking of different scenarios as well as prioritizing among alternative risk scenarios (Campbell et al., 2000).

Especially in ecological systems, limited data and knowledge can hinder modelling efforts, as they constrain it to simpler model structures that involve more assumptions, in these cases the Bayesian network approach can still be applied (Hamilton & Pollino, 2012). Also, Bayesian networks can be developed as casual models, which can be used to assist risk prioritization to help understand pathways of hazard and vulnerability relations better (Sperotto et al., 2017).

A recent paper by *Carriger & Barron (2020)* showed how Bayesian network estimated the risk quotient by calculating the probability of an exposure distribution exceeding an effect distribution. Their Bayesian network estimated the risk by expanding the standard risk equation to include more uncertainties and variables that influence the risk (Carriger & Barron, 2020). The networks we have created used similar risk quotient calculations though instead on focusing on one terrestrial species, we have included multiple species (e.g. SSD) and tried to carry out a risk characterization for the aquatic environment.

Carriger & Barron (2020) also stated that *“the capabilities for performing diagnostic, mixed, and predictive inference make Bayesian networks especially useful for examining the causal factor that could lead to higher or lower risk outcomes”*. The networks we developed use discretisation of continuous variables and with that lose some of the initial precision and information. Nevertheless, another benefit of using Bayesian networks over other probabilistic methods mentioned is the possibility to use dynamic discretization to enable higher resolution and fewer uncertainties associated with the estimations (Carriger & Barron, 2020).

Furthermore, Verdonck et al. (2005) pointed out that there are some unquantifiable uncertainties such as the choice of distribution, model and extrapolation uncertainties that remain difficult to quantify some of which may be overcome by using different distribution models than the ones used in this study. An alternative to the exposure modelling we have carried out in this study was presented by Wolf and Tollefsen (2021) showing how Bayesian distributional regression models could be used to better include spatiotemporal conditional variances in exposure assessment and still allow for a distributed PEC (Wolf & Tollefsen, 2021). Therefore, there is possibility and need for further development, e.g. to better account for spatial and temporal variation in exposure and inter-vs. intra-species variation in sensitivity in effect assessment. Anyhow, Bayesian networks ability to perform predictive and diagnostic inference (Carriger & Barron, 2020) still enable a good understanding of the network and transparency. Thus, they can offer a transparent way of evaluating the required characterization of uncertainty for pesticide risk assessment as well as for ecological risk assessment in general (Carriger & Newman, 2012).

## 5 Conclusion

This study demonstrates that Bayesian network modelling is a promising tool for probabilistic calculation of a risk quotient, which is commonly used in environmental risk assessment of pesticides and other chemicals. A probabilistic risk quotient is a more informative alternative to the traditional single-value risk quotient, which is often interpreted as a binary outcome. The Bayesian network approach provides more opportunities for interpretation, such as the probability of the risk quotient that exceeds not only 1 but also other specified threshold values. The Bayesian network model presented here can easily be mapped to the main steps of traditional risk characterisation frameworks. The Bayesian network approach can still apply an uncertainty factor to account for additional uncertainties that are not captured by the exposure and effects distributions, corresponding to the assessment factor used in traditional risk assessment. Thus, Bayesian networks can offer a transparent way of evaluating the characterization of uncertainty required for pesticide risk assessment as well as for ecological risk assessment in general (Carriger & Barron, 2020).

Our planned further development of this Bayesian network includes extending the model for cumulative risk assessment of pesticide mixtures in the aquatic ecosystem. Furthermore, we will incorporate climate and agricultural scenarios to predict environmental risk of pesticides under future conditions.

## Contact information

Sophie Mentzel

## Software availability

BN modelling using Netica 6.05 (www.norsys.com)

## Appendix A

**Table A.1.**
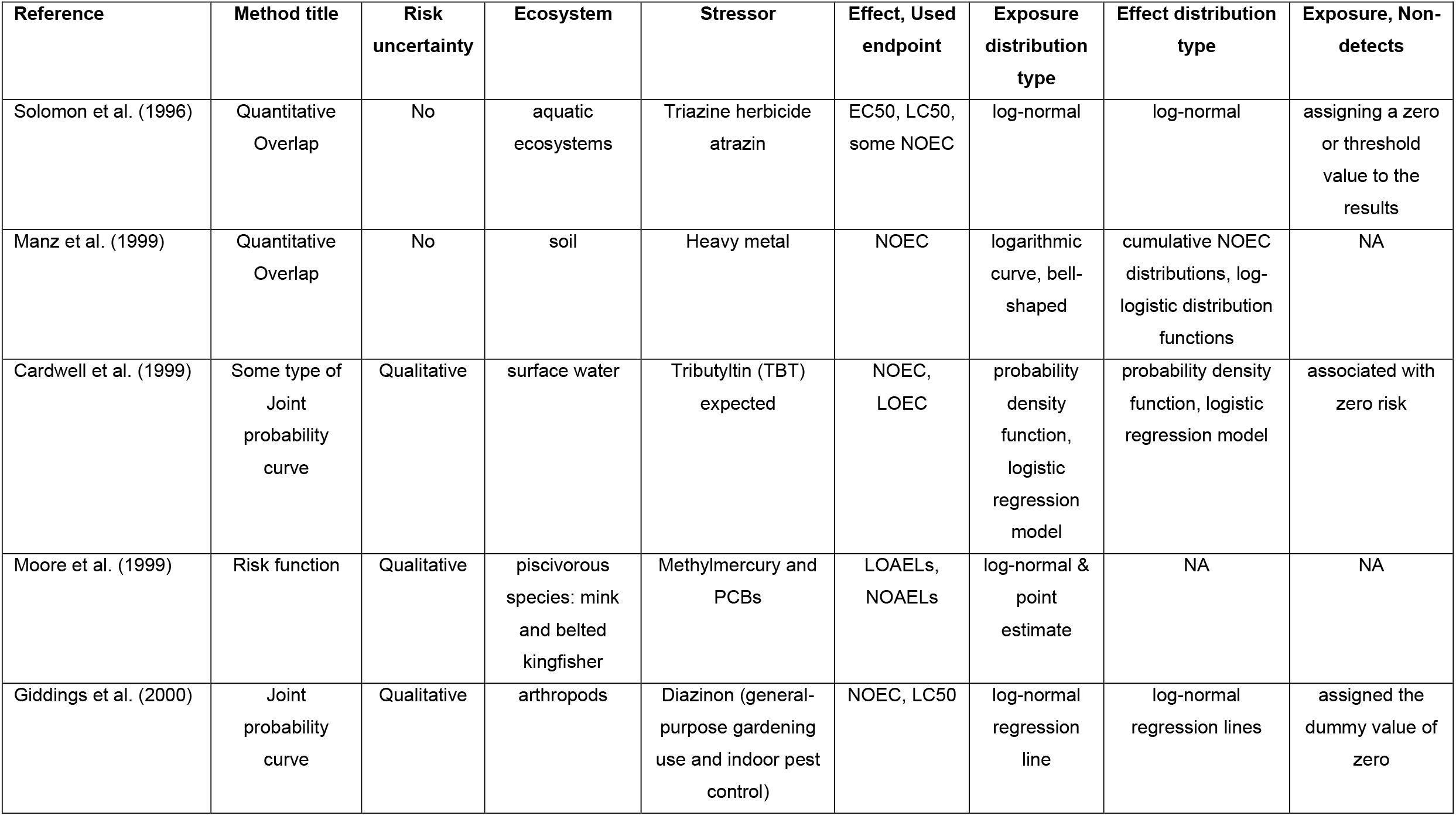
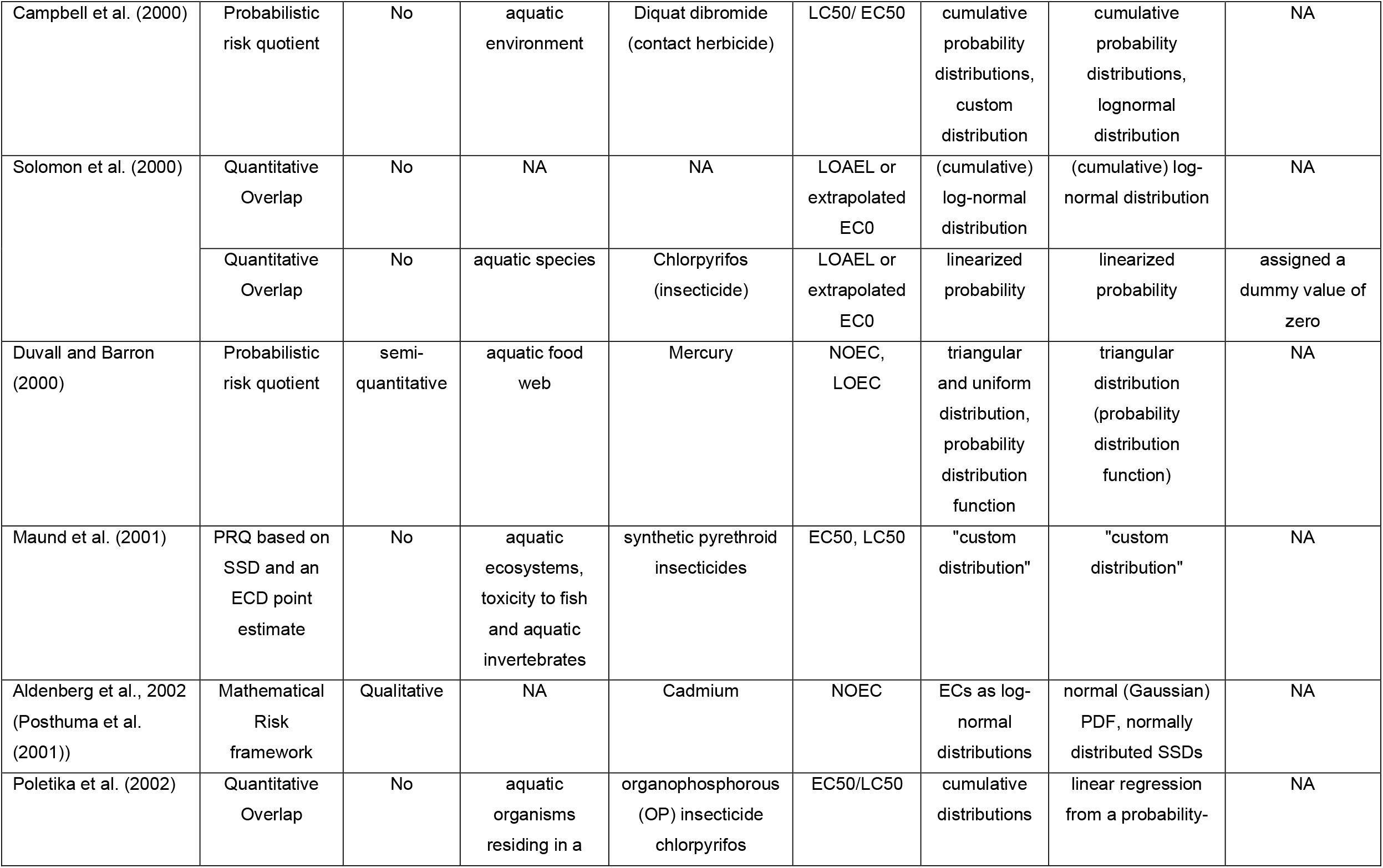
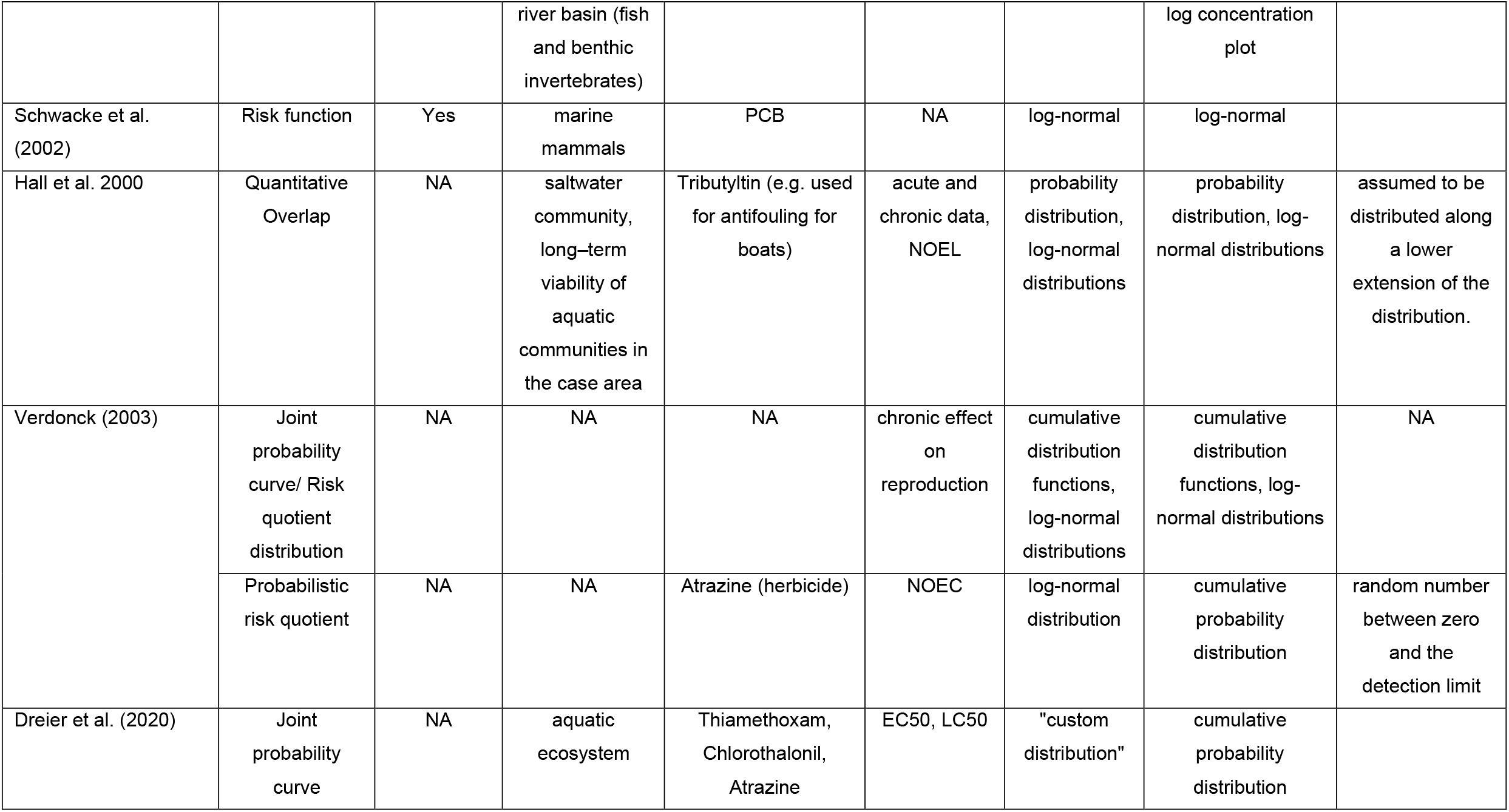
Overview of several probabilistic assessment methods

